# Using Manganese-Enhanced MRI to visualize Magnetogenetic-based Neuromodulation

**DOI:** 10.1101/2025.06.20.660767

**Authors:** Brianna Ricker, Nir Dayan, Galit Pelled, Assaf A. Gilad

## Abstract

**Purpose:** Investigation of the electromagnetic perceptive gene (EPG) protein and garnering evidence to suggest its use as a magnetogenetic tool for neuromodulation. Activation of EPG by electromagnetic field increases intracellular calcium levels, thus, we aimed to determine whether EPG influences the analogous manganese ion. This work yields potential for manganese-enhanced MRI (MEMRI) to be used to monitor EPG acting as a neuromodulator.

**Methods:** HEK293FT cells expressing EPG were exposed to a MnCl_2_ solution and stimulated with a static or electromagnet. Excess MnCl_2_ was washed off the cells, followed by their collection and lysis. T_1_ map measurements of the lysates were obtained to gauge the presence of intracellular Mn^2+^. Several controls were employed to critically evaluate whether EPG can influence Mn^2+^ dynamics.

**Results:** Lysate of cells expressing EPG showed significantly lower T_1_ values compared to cells without EPG that were exposed to the same MnCl_2_ solution and magnetic stimulus.

**Conclusion:** Magnetic activation of EPG increases the uptake of Mn^2+^ into the cell. By influencing ions pertinent to neuronal function, we demonstrate the potential of MEMRI to monitor EPG neuromodulatory activity.

## Introduction

Neuromodulation aims to enhance or inhibit neuronal function by directly altering the activity of neurons, particularly in cases where direct pharmaceutical intervention is not feasible. While deep brain stimulation (DBS) and transcranial magnetic stimulation (TMS) are considered the gold standard, advances in DNA technologies have sparked growing interest in developing techniques that activate specific subpopulations of neurons, preferably remotely, thereby reducing background noise and increasing treatment precision^1^. Indeed, in recent years, several new technologies have emerged, including optogenetics^2,3^, chemogenetics^4^, and magnetogenetics^5,6^.

The electromagnetic perceptive gene (EPG) protein, discovered in the glass catfish (*Kryptopterus vitreolus*), is known to sense and respond to magnetic fields^7,8^. This response has been primarily observed as an influx of calcium ions^7,9–11^. The prospect of using EPG as a means of external control over biological processes is attractive^12,13^ and its synergy with the common signaling molecule Ca^2+^ opens possibilities for remote control of neurological systems^14,15^.

Visualizing and quantifying the levels of magnetogenetics-based neuromodulation and their effect in vivo through MRI are of great interest^16^. Manganese-enhanced MRI (MEMRI) is an imaging technique that uses the paramagnetic properties of manganese ions (Mn^2+^), which enter active neurons through voltage-gated calcium channels. This technique has been particularly paramount for gaining high-resolution data in excitable tissues, such as structures within the brain, due to the high density of voltage-gated calcium channels present^17–20^. This technique works because both calcium and manganese ions possess a similar ionic radii and adopt a 2+ charge. If EPG is able to influence Mn^2+^ as it does Ca^2+^, this posits Mn2+ as a mean to visualize magentogenetics-based neuromodulation via MEMRI.

In addition to using MEMRI to facilitate noninvasive imaging of EPG activity, EPG can improve the specificity of MEMRI. Spatial and temporal expression of EPG in specific subpopulations of neurons in conjunction with MEMRI could further enhance the resolution of neuronal structures or allow MEMRI to visualize details from less excitable tissues that have reduced abilities to actively uptake Mn^2+^. MEMRI has been used in several pre-clinical models of neurological conditions^20–23^, and in brain imaging with manganese-based nanoparticles^24–26^. Thus, a fusion of these technologies could allow MEMRI to significantly accelerate the development of magnetogenetics as a transformative tool for neuromodulation.

## Methods

### Mammalian tissue culture

HEK293FT cells (ATCC) were grown and maintained in Dulbecco’s Modified Eagle Medium (Gibco) supplemented with 10% fetal bovine serum (Gibco), 1% penicillin-streptomycin (Gibco), and 1% geneticin (Gibco). For experiments, cells were seeded at a density of 0.2×10^6^ in 35 mm dishes pre-treated with poly-D-lysine (Gibco). The following day, cells were transfected using the Lipofectamine 3000 kit (Thermo) with the relevant plasmids containing EPG or other proteins.

### Stimulation and lysis

24 hours post-transfection, media was removed from the surface of the cells and a solution of 10mM HEPES + 100mM NaCl ± 100µM MnCl_2_ was placed over the cells. Cells were then exposed to different stimuli from electromagnetic coils^27^ (sham: 14.5A, 0mT; 15s on 5m off for 4 pulses / active: 4.5A, 14.5mT; 15s on 5m off for 4 pulses), two 50mT static magnets, or no stimulus. After the 16-minute incubation period with or without magnetic stimulus, the solution was removed, and the cells were washed once with PBS. The cells were covered with 200μL of 10mM HEPES and detached from the dish using a cell scraper. The suspended cells were collected in a microcentrifuge tube and lysed by several freeze-thaw cycles in liquid nitrogen.

### MRI

T_1_ map measurements of the lysate were taken with a 7 Tesla Bruker MRI, 9 TRs (33, 122, 297, 497, 727, 1577, 2777, 4809, 12496 ms) were measured using a RARE sequence, RARE factor = 1, FOV = 17 × 17 mm^2^, slice thickness = 1.2 mm, matrix size = 64 × 64, spatial resolution = 0.25 × 0.25 mm^2^)^28^. Values for the 10mM HEPES solution were also measured and then subtracted from the T_1_ values of the samples to standardize data between scans.

## Results

Electromagnetic stimulus causes an influx of Mn^2+^ facilitated by EPG As shown in Figure 1A, we observe a decrease in T_1_ values for all groups that were exposed to MnCl_2_. However, the group expressing EPG that was exposed to active electromagnetic stimulus has a statistically significant decrease in T_1_ compared to cells without EPG and stimulus (p-value 0.0287). In Figure 1B, we compare two additional proteins to EPG. CD59 is a human homolog of EPG that is similar in size and structure that does not respond to magnetic fields. EPG(B.g.) is an engineered version of EPG based on a homologous protein from the fish *Brachyhypopomus gauderio* that increases calcium concentration in mammalian cells faster, but to the same magnitude as EPG ^11^. We observe that CD59 did not produce any significant differences in T_1_, whereas the EPG(B.g.) exposed to the active electromagnet stimulus has a significantly decreased T_1_ compared to the relevant control group (p-value 0.0030).

**Figure 1.**
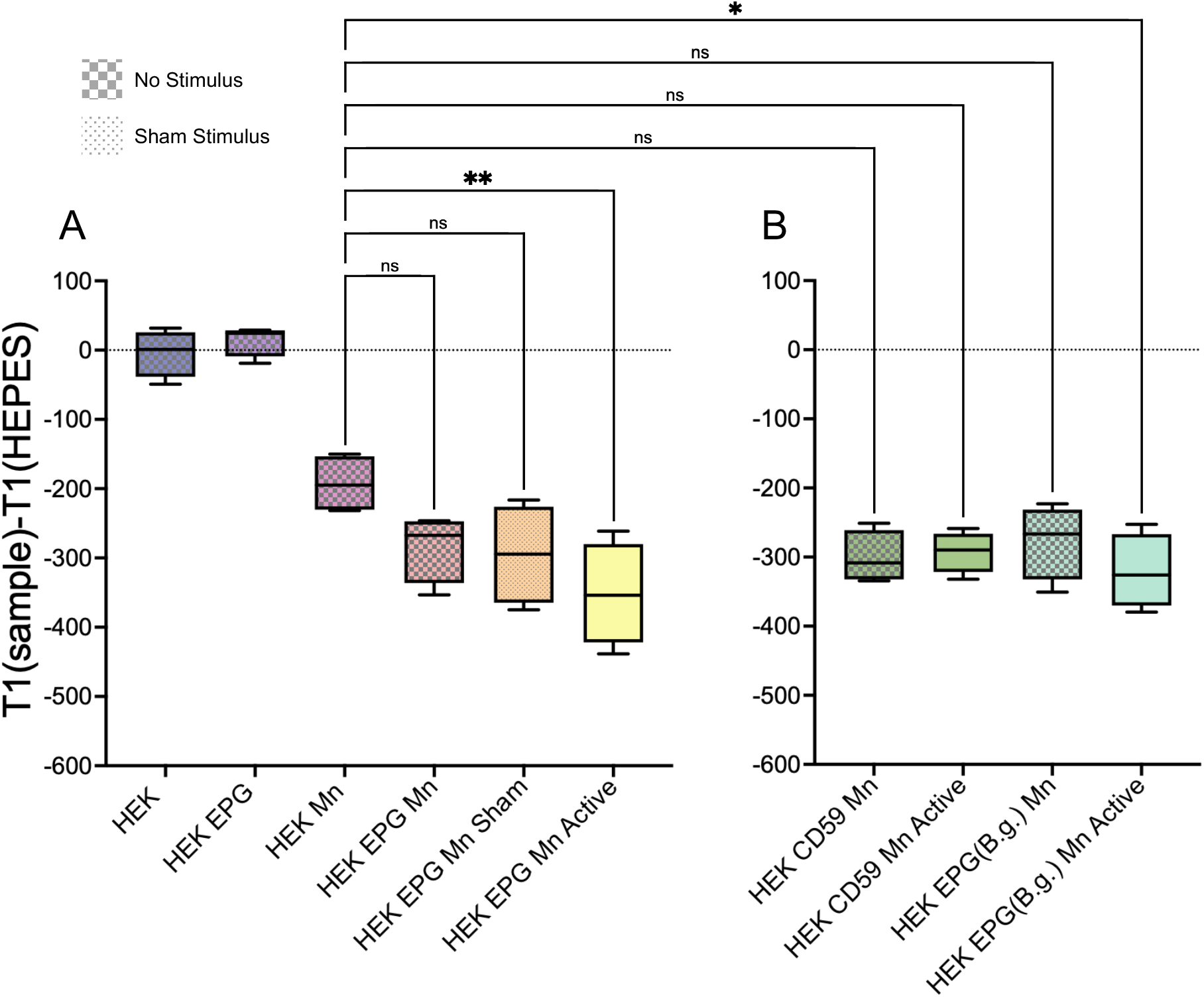
Mn2+ uptake facilitated by electromagnetic coil stimulation. Lysate of HEK293FT cells (HEK) exposed to various conditions (i.e. transfected with different plasmids, exposed to MnCl_2_ (Mn), exposed to stimuli) and subject to MRI. **A.** T_1_ values for lysate of cells with or without EPG and Mn under no stimulus (checked boxes), sham stimulus, or active stimulus. **B.** T_1_ values for lysate of cells expressing control proteins with and without active magnet stimulus. All groups are representative of four experiments (n=4). Ordinary one-way ANOVA was used to determine significance between groups.

Static magnet stimulation also facilitates Mn^2+^ influx via EPG In Figure 2A, while a decrease in T_1_ values was observed in all groups exposed to MnCl_2_, a significant decrease in T_1_ was observed only for those cells expressing EPG that received static magnet stimulus (p-value 0.0005). We also observe significant differences between groups that received EPG versus the counterpart that did not receive EPG (p-values <0.05).

**Figure 2.**
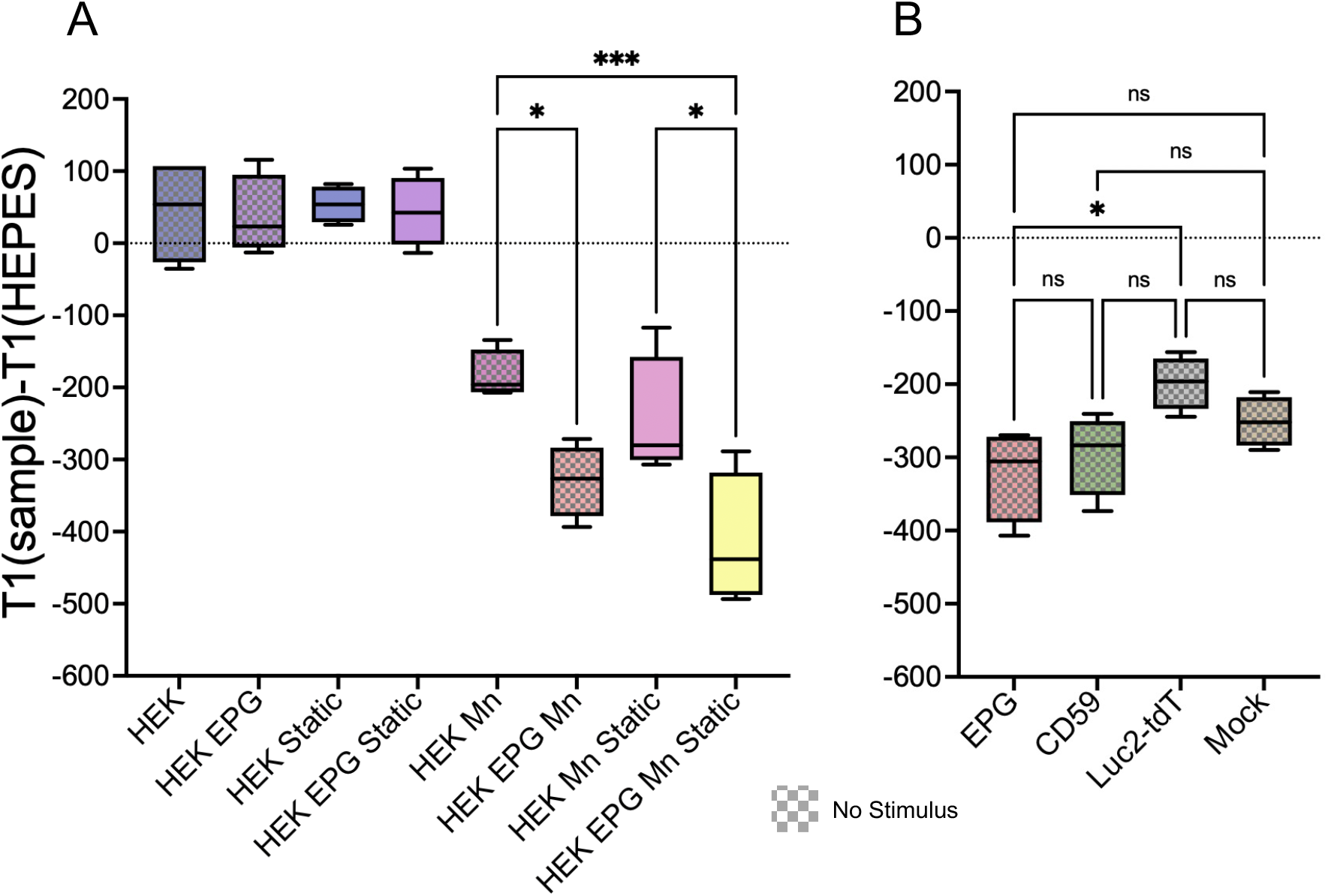
Mn2+ uptake facilitated by static magnet stimulation. T_1_ values for lysate of HEK293FT cells (HEK) exposed to various conditions (i.e. transfected with different plasmids, exposed to MnCl_2_ (Mn), exposed to static magnets (Static)) and subject to MRI. **A.** T_1_ values for lysate of cells with or without EPG and Mn under no stimulus (checked boxes), and static magnet stimulus. **B.** T_1_ values for lysate of cells expressing different plasmids exposed to MnCl_2_.

This prompted us to determine if transfection has an impact on the T_1_ values observed.

Figure 2B shows HEK cells transfected with EPG and several control plasmids: CD59 (GPI anchored; as is EPG), Luc2-tdT (cytosolic), and mock transfected. The general trends suggest that transfection of a membrane-associated proteins (EPG and CD59) results in a lower T_1_ value; however, only the combination of EPG with Mn shows a statistically significant reduction in T_1_ (p-value 0.0193).

All groups are representative of four experiments (n=4). Ordinary one-way ANOVA was used to determine the significance between groups.

## Discussion

This work represents progress in using MEMRI for imaging in conjunction with genetically encoded tools. Bartelle, Turnbull, and colleagues engineered the divalent metal transporter 1 (DMT1) and used MEMRI to characterize DMT1-expressing cells, both in vitro and in vivo in mouse models of the developing brain^29^. Recently, Rallapalli, Koretsky, and colleagues have shown that the manganese ion transporter Zip14 is visible in vivo using MEMRI. The viral delivery-induced neuron-specific overexpression of Zip14 resulted in significant enhancement MRI contrast^30^.

The results presented here indicate that EPG is able to induce Mn^2+^ influx in response to magnetic stimulation. The current data shows small, but significant differences in Mn^2+^ based contrast, which will open the possibilities for EPG to be engineered and improved upon. In its current state, EPG has influence over at least Ca^2+^ and Mn^2+^ which likely compete with each other for space within the osmolarity of the cell, leading to small changes like those we have observed. One future improvement to EPG would be introducing a specificity mechanism for Ca^2+^ and Mn^2+^ for more significant changes and robust use.

One possible explanation for the lower T_1_ in cells transfected with EPG and CD59 is that the overexpression due to the CMV promoter and transient transfection of these proteins and their resultant insertion into the membrane disrupts the native membrane dynamics and makes the cells ‘leaky’ to Mn^2+^.

These in vitro experiments also yield potential for experimentation in vivo, particularly in excitable tissue like the nervous or cardiac systems, where the appropriate machinery for influencing Mn^2+^ dynamics is more accessible to EPG than in a HEK cell. These in vivo models can be used for studying neurological disorders and developing new treatments based on magnetogenetics and neuromodulation. Using EPG in conjunction with MEMRI yields potential for studying magnetogenetic neuromodulation, and potential for improved Mn^2+^ based MRI contrast.

## Conclusion

This newfound property of EPG to influence Mn^2+^ dynamics within biological systems expands its capability to be engineered as a magnetogenetic therapeutic. Its use with MEMRI can facilitate magnetogenetics toward clinical translation for neural manipulation, thereby enhancing options for treating neurological disorders. Moreover, this data shows that EPG can be further developed as a part of new magnetogenetic technologies.

## Acknowledgements

A.A.G acknowledges financial support from the NIH/NIBIB: R01-EB031008; R01-EB030565; R01-EB031936.

